# Telomere length and cognitive function among middle-aged and older participants from communities underrepresented in aging research: A preliminary study

**DOI:** 10.1101/2024.10.14.618331

**Authors:** Lauren W. Y. McLester-Davis, Derek Norton, Ligia A. Papale, Taryn T. James, Hector Salazar, Sanjay Asthana, Sterling C. Johnson, Diane C. Gooding, Trevor R. Roy, Reid S. Alisch, Stacy S. Drury, Carey E. Gleason, Megan Zuelsdorff

## Abstract

**Objective:** Accelerated biological aging is a plausible and modifiable determinant of dementia burden facing minoritized communities, but is not well-studied in these historically underrepresented populations. Our objective was to preliminarily characterize relationships between telomere length and cognitive health among American Indian/Alaska Native (AI/AN) and Black/African American (B/AA) middle-aged and older adults.

**Methods:** This study included data on telomere length and cognitive test performance from 187 participants, enrolled in one of two community-based cognitive aging cohorts and who identified their primary race as AI/AN or B/AA.

**Results:** Nested multivariable regression models revealed preliminary evidence for associations between telomere length and cognitive performance, and these associations were partially independent of chronological age.

**Discussion:** Small sample size limited estimate precision, however, findings suggest future work on telomere length and cognitive health in underrepresented populations at high risk for dementia is feasible and valuable as a foundation for social and behavioral intervention research.

## Introduction

As the U.S. population ages, the incidence and prevalence of Alzheimer’s disease and related dementias (ADRD) is growing, creating substantial financial burden for individuals, families, and communities (*Alzheimer’s Association 2024 Alzheimer’s Disease Facts and Figures*, 2024). The same is true for subclinical cognitive impairment, which represents a risk factor for future progression to dementia, and even in a pre-diagnostic stage presents other risks for older adults: age-related cognitive changes well below clinical thresholds robustly predict incident falls, loss of independence, and even mortality (Aliberti et al., 2019). Importantly, the burden of ADRD is not equitably distributed across populations; social disparities are prominent. Minoritized populations, including American Indians/Alaska Natives (AI/AN) and Black/African Americans (B/AA), experience disproportionately high risk of cognitive impairment and dementia relative to non-Hispanic white peers (Mayeda et al., 2016).

Given the impact of cognitive aging on quality of life, and a continued lack of accessible disease-modifying therapies, preventing incident cognitive impairment is imperative. Fortunately, a substantial proportion of ADRD risk is modifiable under conditions that support cognitive health. A recently updated meta-analysis found that fourteen well-studied, primarily behavioral and biomedical risk factors account for 45% of global dementia; that proportion is even greater in racially minoritized populations (Livingston et al., 2024; Lee et al., 2022). Advancing age is considered one of the few non-modifiable determinants of ADRD-related impairment risk, and certainly this is the case for *chronological* age. However, the correlation between chronological age and *biological* age varies across individuals and populations, where biological age is influenced by disadvantageous environmental, socioeconomic, behavioral, and biomedical exposures that create wear and tear on biological systems (Kennedy et al., 2014). When one’s biological age outpaces one’s chronological age, the aging process is accelerated, and so is the time to cognitive impairment (Movérare-Skrtic et al., 2012). This distinction is crucial for ADRD prevention research: biological aging is strongly predictive of functional declines characteristic of later life, and is modifiable not only through addressing the exposures previously mentioned but also targeted intervention (Wu et al., 2021).

The measurement of telomere length (TL) is increasingly used as an easily obtained marker of biological age and has significant potential to support translatable mechanism-focused ADRD science (Vaiserman & Krasnienkov, 2020). Telomeres are the protective DNA and protein structures at the ends of human chromosomes that maintain genomic stability. Measurement of TL may be assessed from the variety of tissue samples commonly collected in aging-focused cohort studies. The shortening of telomeres is an intrinsic part of aging, but rates of telomere attrition are highly heterogeneous and even short-term behavioral interventions targeting changes in TL have shown promise (Buttet et al., 2022). Further, multiple lines of research support measures of TL as a predictor of cognitive outcomes, including preclinical changes that herald progression to ADRD (Byun et al., 2020), mild cognitive impairment (MCI), and dementia (Insel et al., 2012; Grodstein et al., 2008; Kume et al., 2012; Hochstrasser et al., 2012; Liu et al., 2016; Forero et al., 2016; Scarabino et al., 2017; Scarabino et al., 2020). That said, differences in magnitude and even direction of effect for biological age and cognition have been observed across demographic characteristics including race/ethnicity (Avila-Rieger et al., 2022), indicating a crucial need for inclusive methodologies in this discipline.

There is a particularly pressing need for additional research on biological age as a predictor of cognitive risk that prioritizes inclusion of AI/AN and B/AA populations (Thomas et al., 2008; Lukens et al., 2009; Movérare-Skrtic et al., 2012; Mahoney et al., 2019). Racism is a social determinant of inequity, producing systematized differentials in risk exposures across the life course, as well as in the resources to mitigate them (Adkins-Jackson et al., 2023; Glymour & Manly, 2008). It is thus not surprising that advanced biological age compared to chronological age, conceptualized in health equity models as socially-rooted *biological weathering*, has been reported in racially minoritized groups such as the B/AA population in the United States (Geronimus, 1992; Elliott et al., 2021). However, despite the increased risk for developing ADRD compared to non-Hispanic whites (Mayeda et al., 2016) and reported accelerated biological aging (Brown et al., 2017), AI/AN and B/AA are underrepresented in ADRD research cohorts. The gap in ADRD research for AI/AN and B/AA includes both inadequate representation in both observational studies and clinical trials, and arises as a result of burdensome and/or invasive protocols as well as inadequate strategies for culturally-responsive educational outreach and engagement (Olson & Albensi, 2020; Gilmore-Bykovskyi et al., 2019). Recruitment science studies identify lack of access to research centers and distrust of investigators as barriers to participation in research among racially minoritized communities, who are also less likely to consent to studies examining genetic data (Leibel et al., 2020). Comprehensive inclusive science programming addresses underrepresentation as an institutional problem, and pairs outreach with both material and emotional support for accessible participation and cultural safety (Gilmore-Bykovskyi et al., 2022; Gleason et al., 2019).

Strong academic-community partnerships built across time, and study leadership by an Indigenous investigator who is well-known to local Tribal and Black communities, created a unique opportunity for our team to improve understanding of biological age and cognitive health in two populations historically excluded from and underserved by ADRD research. With approval and buy-in from a Tribal and a B/AA Community Advisory Board, this study provides preliminarily examination of TL as a predictor of cognitive function in a sample of middle-aged and older AI/AN and B/AA adults.

## Methods

### Source Studies, Community Engagement, and Participants

Data were collected from participants enrolled in two longitudinal cohort studies at the University of Wisconsin: 1) the Wisconsin Registry for Alzheimer’s Prevention (WRAP) study and 2) the University of Wisconsin Alzheimer’s Disease Research Center (ADRC) clinical core (Johnson et al., 2018; Allison et al., 2021). All B/AA participants were co-enrolled in the ancillary African Americans Fighting Alzheimer’s in Midlife (AA-FAIM) study. The WRAP is a longitudinal observational study, with rolling recruitment and enrollment, of middle-aged adults who are cognitively normal at study entry. Approximately 70% of the cohort has a self-reported parental history of ADRD (Clark et al., 2018). The University of Wisconsin ADRC clinical core is another longitudinal observational study, and is primarily composed of participants without impairment, but also enrolls participants with MCI and dementia and retains participants who develop impairment during follow-up. WRAP and ADRC participants complete comprehensive in-person visits, annually or biennially, that include neuropsychological cognitive exams, surveys of self-reported health and behavioral data, a physical exam by a nurse practitioner, and collection of blood for clinical and laboratory assays.

For B/AA participants, the AA-FAIM study provides support for community engagement while leveraging the data infrastructure of the WRAP and ADRC clinical core studies. Wisconsin ADRC funding likewise supports outreach and engagement for local AI/AN communities. All source studies ultimately dedicate substantial resources including staff effort to longstanding community partnerships, engaging a multifaceted approach to inclusion to grow a diverse cohort of highly engaged participants. Specific to the present investigation, this approach included multiple community-based talks on biological aging, provided by a researcher with Indigenous lived experience and dual expertise in neuroscience and ethics of data sovereignty (LWYMD). These presentations have generated substantial community and participant interest in our team’s work on TL as a cognitive impairment risk factor.

Given project goals of assessing biological age among middle-aged and older adults from populations that are underrepresented in, and therefore underserved by, most ADRD cohorts, this study sample ultimately included 187 participants who self-identified racially as B/AA or AI/AN, provided whole blood, and had concurrent complete cognitive and covariate data available for at least one study visit. The present study was approved by and has received ongoing feedback from the ADRC’s Tribal and B/AA Community Advisory Boards. The University of Wisconsin Institutional Review Board approved all study procedures, each participant provided signed informed consent before participation, and all research was completed in accordance with the Helsinki Declaration as previously described (Johnson et al., 2018). Importantly, we also obtained community approvals for the research direction and these analyses. Formal Tribal approvals were obtained for the research teams to be on Tribal lands to recruit participants. Additional community input for the analyses themselves were provided by community advisors. For our work with Indigenous participants, the Community Advisory Board’s role and function were approved by the Tribal government. TL was measured at Tulane University and Tulane University’s Institutional Review Board determined TL measurement for these de-identified DNA samples as exempt research.

### DNA Extraction and Telomere Measurement

Whole blood samples were collected in 10mL EDTA tubes, mixed by rocking for 5 minutes, and aliquoted into 5mL tubes before storage at −80°C. Samples were thawed and genomic DNA was extracted at the University of Wisconsin using the Gentra Puregene Blood Kit following the manufacturer’s protocol (Qiagen). Approximately 200ng of DNA per sample was aliquoted, labeled with a de-identified subject ID, and shipped to Tulane University on dry ice overnight for DNA quality assessment and TL analysis.

Integrity and purity of genomic DNA samples were assessed via BioTek Epoch Microplate Spectrophotometer and with the Invitrogen Qubit dsDNA Broad Range Assay Kit. The average 260/280 ratio for all DNA samples was 1.91, the average 260/230 ratio was 0.99, and the average double-stranded DNA concentration was 149.16 nanogram per microliter (ng/µl). The average relative TL was determined from the telomere repeat copy number to single gene (albumin) copy number (T/S) ratio using an adapted monochrome multiplex quantitate real-time polymerase chain reaction (MMQ-PCR) via a BioRad CFX96 as previously described (Cawthon, 2009; Drury et al., 2014). Samples were assayed in triplicate with different well positions on a duplicate plate, using a 7-point standard curve from a peripheral white blood cell DNA standard ranging from 0.0313ng to 2ng, with an average of 1.25 replicates of the standard removed per plate. The average efficiencies of telomere and single copy gene primers were 94.34% and 95.52% respectively, with an average R^2^ of 0.99 for telomere and single copy gene standard replicates. All efficiencies were between 90-110% and within 10% of each other to eliminate plate to plate variability. The average slope and y-intercept for the telomere standard curve were −3.47 and 4.93, respectively. The average slope and y-intercept for the single copy gene standard curve were −3.44 and 12.88, respectively. Coefficients of variations were 4.10% for within triplicate variation on average and 1.90% for between duplicate plate variation on average. Only 13% of samples were repeated once for a minimum of 5 replicates with passing quality control criteria. The intra-class correlation (ICC) for all samples passing quality control criteria was calculated as 0.887 (CI: 0.862, 0.908) in accordance with previously described computations (Eisenberg et al., 2020).

### Neuropsychological Assessments

Performance on tests of executive function and episodic memory, cognitive domains sensitive to early age-related change, served as key outcomes (Lezak, 2004). The former was assessed with Trail Making Test A (TMT A) and B (TMT B). TMT A and B consist of 25 circles on paper containing numbers or numbers and letters. Participants were instructed to connect the numbers as quickly as possible in ascending order in TMT A, and in an ascending pattern alternating numbers and letters in TMT B. The TMT A is a visual-motor attentional task, while the TMT B is a visual-motor executive functioning task, with additional demands on working memory and cognitive switching. Performance was measured in the time to complete the test; errors were corrected as they occur and were included in the total time-to-completion (Corrigan & Hinkeldey, 1987). In the WRAP and ADRC clinical core protocols, the normative time 300-second cutoff for the TMT B was extended to 600 seconds to allow for greater variability in task performance (Strauss et al., 2006).

Episodic memory was assessed with the well-known and validated Rey Auditory Verbal Learning Test (RAVLT; *Rey Auditory Verbal Learning Test ^TM^*, 1996). The RAVLT requires participants to learn a list of 15 unrelated words and immediately recall as many as possible, over 5 Learning Trials. Each correctly recalled word counts for one point. Summing the words recalled in each trial provides a score representing immediate memory (RAVLT Sum of Learning Trials 1-5). Following a delay of ∼30 minutes and the reading of a “distractor list” of 15 new words, participants were asked to recall as many of the original words as possible (RAVLT Long Delay) for a measure of delayed recall.

### Self-Identified Race and Other Covariates

WRAP and ADRC participants were asked to self-identify their race/ethnicity at their baseline study visit; formatting at the time of enrollment for participants in the current analysis required participants to select a single response for “primary race” with optional follow-up questions on “secondary” and “tertiary” race. A separate question asked participants to identify ethnicity as Hispanic/Latino or non-Hispanic/Latino. As noted, selection into the current study sample was based on primary, secondary, or tertiary race self-reported as “American Indian/Alaska Native” or “Black/African American.” While sampling was based on ensuring representation of racially minoritized communities generally, racialization uniquely shapes lived experiences and thus both biological age and cognitive performance. Though self-identified race is only modestly correlated with ancestry at a population level, there is also some evidence for ancestry-related racial differences in TL (Hansen et al., 2016). Accordingly, participants were also aggregated into three racial groups to model race as a covariate: primary race as AI/AN with no secondary racial group identified, primary race as B/AA with no secondary racial group identified, and multiracial.

Other covariates were likewise selected *a priori* based on empirical evidence for their correlations with both the exposure and outcomes of interest. In addition to self-identified race, these included chronological age at visit, analyzed as a continuous variable; educational attainment in years, also analyzed as a continuous variable and Winsorized at 20 years if >20 years; and sex, which at the time of sample enrollment required a binary response (“Male” or “Female”) and was analyzed as a categorical variable in the current study.

### Data Analyses

Comparisons of sample characteristics for AI/AN, B/AA, and multiracial participants were assessed with Kruskal-Wallis tests for continuous variables, and Fisher’s exact test for categorical variables. To assess relationships between TL and cognitive performance, we used standard multivariable linear regression. Previous studies including international meta-analyses have found associations to depend on covariate selection for a given model (Zhan et al., 2018). Thus, covariates were incrementally added to a series of nested models to assess the contributions of key potential confounders to observable relationships: (1) a base model controlling for self-identified sex and educational attainment, (2) a model adding self-identified race, and (3) a model adding chronological age. Separate models were run with each of the four cognitive tests as a different dependent variable (TMT A Time and B Time, RAVLT Sum of Learning Trials and Long Delay). For the TMT outcomes, these values were natural log transformed before model fitting to avoid issues with heteroskedasticity. All fitted models were checked for conformance to regression assumptions. There were no major concerns for residual trends, heteroskedasticity, non-normality, and no overly influential data points were noted.

In order to delineate contributions of biological and chronological age, we formally assessed change in the estimated TL regression coefficient when chronological age was included in the model (i.e., from model structure 2 to model structure 3). We utilized non-parametric bootstrapping of the fitted data, collecting the coefficient change over 2000 iterations, and constructing 95% confidence intervals (CIs) on the change using the BCa method.

As noted, the current preliminary study and dissemination of its results was originally motivated by community interest in the research question and a desire to see more inclusive work on TL and cognitive aging. To assist with interpretation of findings and facilitate future work in this area, we also utilize outcomes from these models to conduct a power analysis. Post-hoc power analysis of the TL effect in model structure 3 was assessed using the fitted data (TL and outcome variance) and the fitted model estimates (estimated TL coefficient, R^2^, and number of coefficients in the model). This was used to generate sample size for 80% power to detect the estimated TL effect if more data were available in our cohorts or others in future research.

All analyses were performed in R version 4.4.0 (R Core Team, 2024) using the *boot* package (Canty & Ripley, 1999; Davison & Hinkley, 1997) for bootstrapping, and the *pwrss* package (Bulus, 2022) for the post-hoc power calculations. Statistical significance was assessed at the 5% level for all analyses.

## Results

### Participants Demographics, TMT and RAVLT Scores, and TL Distribution

Full sample characteristics are provided in Table 1. Briefly, the sample (N=187) was majority (74.9%) with an average age of 58.60 years (SD = 8.20 years). 8.6% of participants reported more than one racial and ethnic identity, with 12.8% of the sample reporting AI/AN only and the remaining 78.6% reporting B/AA only. In the full sample, participants reported an average of 14.60 years (SD = 2.32 years) of formal educational attainment. The average time of participants for completing the TMT A was 32.90 seconds (SD = 12.36 seconds) and for the TMT B the average time to completion was 99.00 seconds (SD = 56.20 seconds). These average times are within limits per normative data (Lezak, 2004). The average number of words recalled across RAVLT Sum of Learning Trials 1-5 was 44.19 (SD = 9.02). The average number of words recalled for the RAVLT Long Delay was 8.40 (SD = 3.09). These values are also considered typical of cognitively healthy individuals (Lezak, 2004). There were modest differences in age at visit and TMT B scores across racial groups. There were no significant differences between racial groups for overall mean T/S ratio. The distribution of TL for the sample resembled a normal Gaussian distribution within each racial category as seen in **Figure 1**.

**Figure 1.**
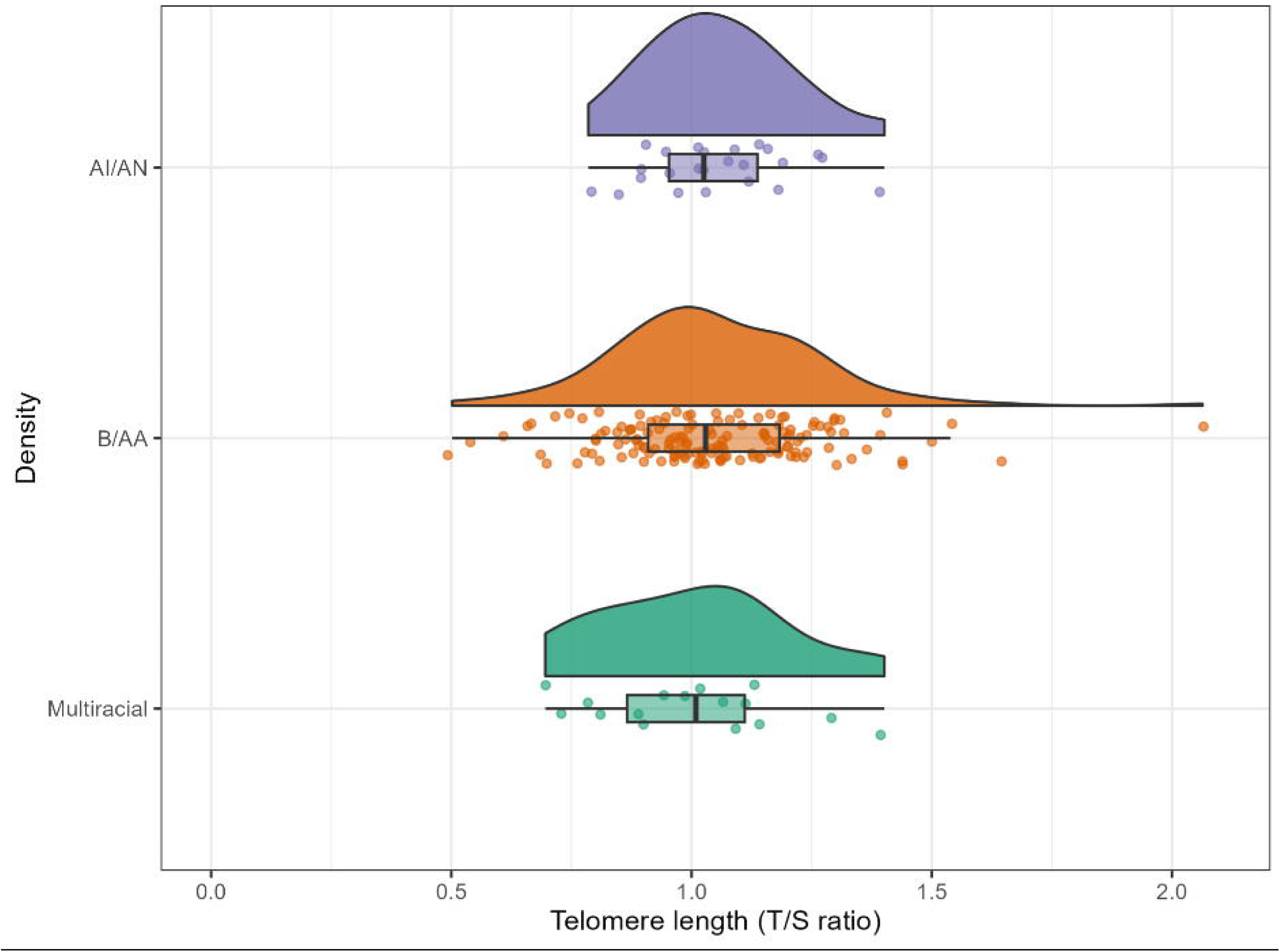
Distribution of telomere length by racial category. Telomere length (TL), measured by telomere to single copy gene (T/S) ratio, is on the x-axis, and the American Indian/Alaska Native (AI/AN) TL measurements are in purple at the top, the Black/African American (B/AA) TL measurements are in orange in the middle, and the Multiracial TL measurements are in green at the bottom.

**Table 1.** Participant characteristics, full sample and by self-identified race.

| Variable | Full Sample | American Indian/Alaska Native | Black/African American | Multiracial | Group Comparison (*p*) |
| --- | --- | --- | --- | --- | --- |
| N | 187 | 24 | 147 | 16 |  |
| Age at visit, years (mean (SD)) | 58.60 (8.20) | 61.97 (7.98) | 57.81 (8.18) | 60.74 | 0.05 |
| Male (N (%)) | 47 (25.1) | 7 (29.2) | 39 (26.5) | 1 (6.2) | 0.18 |
| Formal education, years (mean (SD)) | 14.63 (2.32) | 14.33 (2.06) | 14.61 (2.37) | 15.31 (2.27) | 0.48 |
| Telomere length, T/S ratio (mean (SD)) | 1.04 (0.21) | 1.05 (0.15) | 1.05 (0.21) | 1.00 (0.20) | 0.64 |
| Trail Making Test A, seconds (mean (SD)) | 32.90 (12.36) | 28.62 (9.22) | 33.97 (12.85) | 29.33 (9.99) | 0.06 |
| Trail Making Test B, seconds (mean (SD)) | 99.00 (56.20) | 80.67 (40.55) | 104.27 (59.58) | 76.67 (24.90) | 0.01 |
| RAVLT Sum of Learning Trials 1-5, summed words (mean (SD)) | 44.19 (9.02) | 43.42 (5.98) | 44.29 (9.07) | 44.53 (12.53) | 0.70 |
| RAVLT Long Delay, words recalled (mean (SD)) | 8.40 (3.09) | 8.00 (2.86) | 8.44 (3.08) | 8.67 (3.72) | 0.69 |

### Associations of TL and Cognitive Test Performance

Multivariable linear regression was used via a series of nested models to test whether TL significantly associated with performance on the four cognitive tests when adjusting sequentially for sex, formal educational attainment, race, and chronological age (Table 2).

**Table 2.**
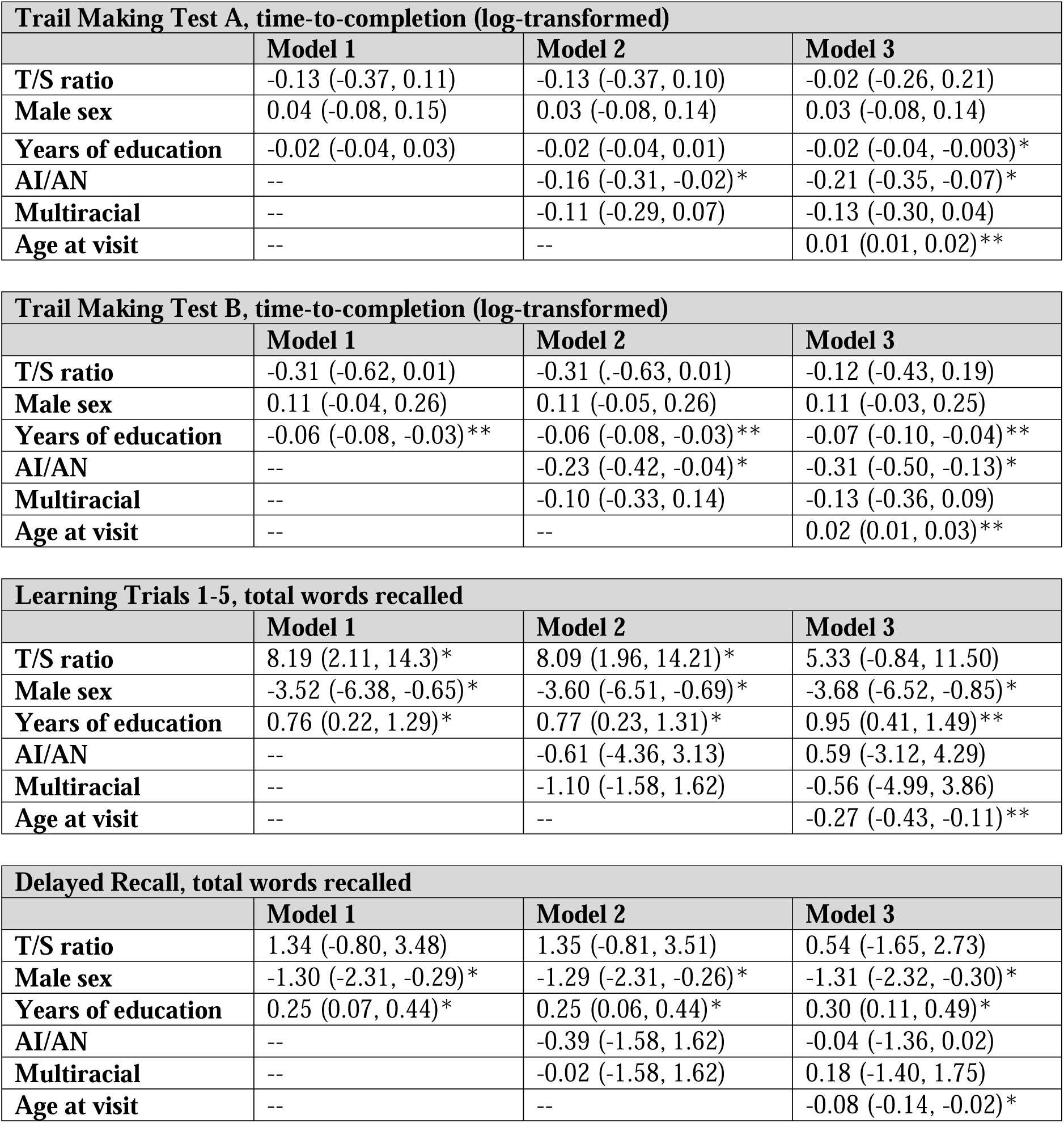
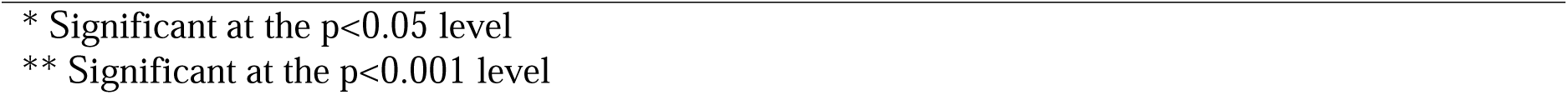
Regression coefficients (95% confidence intervals) for telomere length and cognitive test performance by outcome, race, and model.

There was no evidence for an association between TL and our measure of processing speed, TMT A time-to-completion; the null association did not change between models 1 and 2. Further, when chronological age was added as a covariate in model 3, the estimate for TL was attenuated by nearly 85% (change of 0.11 from −0.133; 95% BCa CI for change: 0.04 to 0.23). Utilizing model 1, the relationship between longer TL and better performance on a measure of more complex executive functioning, TMT B time-to-completion, approached through did not reach statistical significance (*b* = −0.31, 95% CI: −0.62 to 0.01). There was no change in TL estimate in model 2. However, as expected, chronological age confounded relationships between TL and TMT B performance, reducing the estimated Model 2 effect by 62% in magnitude, with a change of 0.12 (95% BCa CI for change: 0.04 to 0.36) in the TL coefficient going from model 2 to model 3.

In terms of measures of episodic memory, TL positively associated with performance on a test of immediate memory and learning, RAVLT Sum of Learning Trials 1-5 (*b* = 8.19, 95% CI: 2.11 to 14.26). This relationship was not substantially attenuated by adjusting for race in model 2, however chronological age did again partially confound the observed association. In model 3, inclusion of chronological age attenuated the TL effect to the point of non-significance as a result (*b* = 5.33, 95% CI: −0.84 to 11.50). The estimated TL effect was ultimately reduced by 34% in magnitude, with a change of −2.75 (95% BCa CI for change: −5.67 to −1.01) in the estimated TL coefficient going from model 2 to model 3. The relationship between TL and the other measure of memory, RAVLT Long Delay, did not reach statistical significance and adjusting for race in model 2 did not change this finding. For RAVLT Long Delay, model 3’s inclusion of chronological age attenuated the estimated TL effect by approximately 60% (estimated change of −0.81 from 1.35; 95% BCa CI for change: −1.82 to −0.24). Overall models of a given TL by performance on each cognitive test are plotted given female participant sex, the average educational attainment of 9.93 years, and the average chronological age of 58.6 years in **Figure 2**.

**Figure 2.**
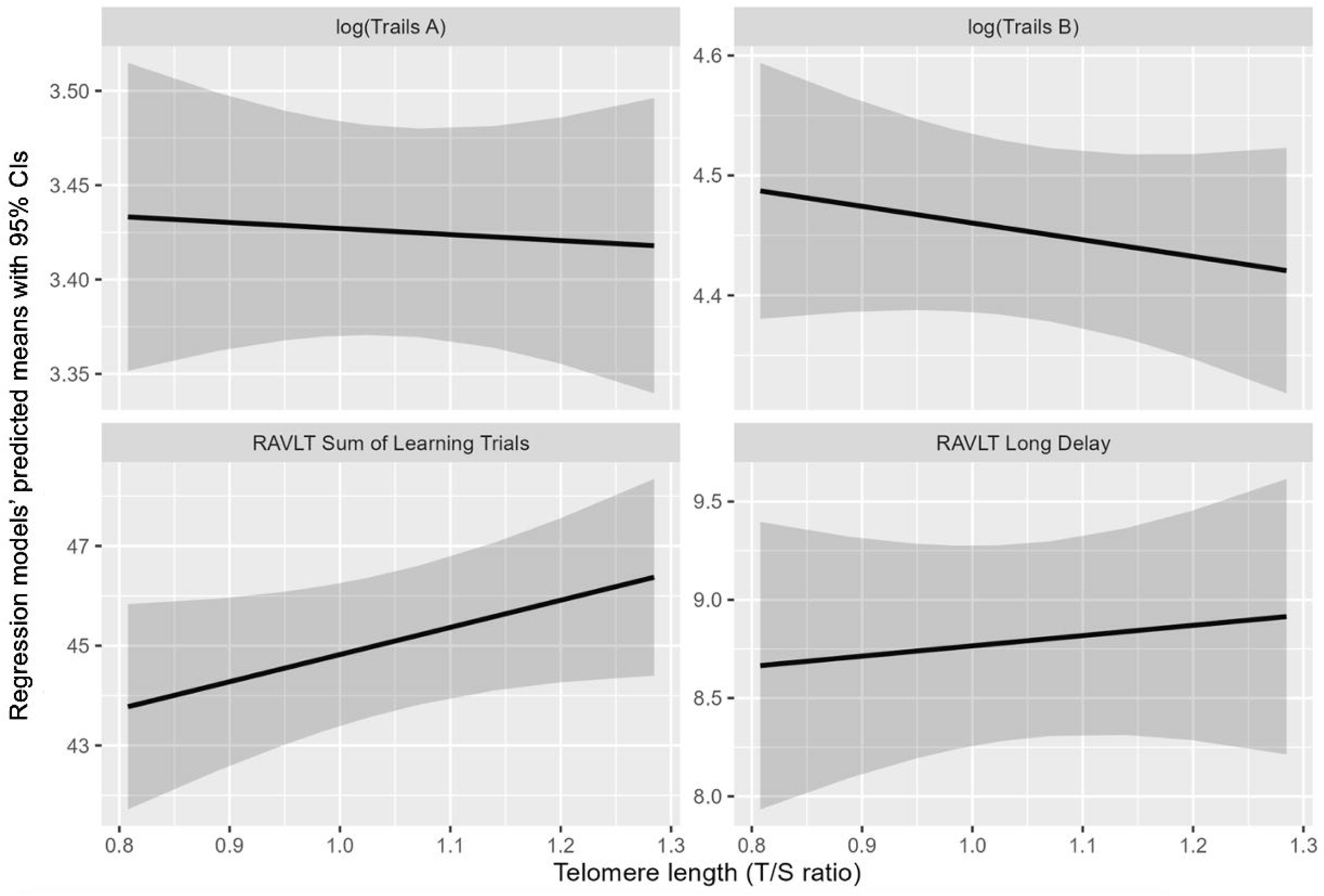
Regression model-predicted means for performance on four cognitive tests for a given telomere length (TL). Regression models have means in dark line and 95% confidence intervals (CIs) in shaded areas by cognitive test. All models are referenced for female, Black/African American participants at the mean participant age of 58.6 years. Scatter plots with linear fit lines of TL with the Trail Making Test (TMT) A time logged (upper left), the TMT B time logged (upper right), the Rey Auditory Verbal Learning Test (RAVLT) Sum of Learning Trials 1-5 (lower left), and the RAVLT Long Delay (lower right). TL was measured by telomere to single copy gene (T/S) ratio.

### Post-Hoc Power Analysis: Predicting Sample Sizes Needed for Future Work

Post-hoc power analysis of the TL regression coefficient significance was conducted using the model 3 structures, fits, and data. For the RAVLT Sum of Learning Trials 1-5, approximately double the number of participants would be needed for 80% power, with an ideal sample size of 401, compared to the current sample size of 187. For other cognitive outcomes with weaker adjusted associations between TL and cognitive function observed, much larger sample sizes would be needed for achieving 80%, with >1500 needed for TMT B, >3600 for RAVLT Long Delay, and >17000 for TMT A.

## Discussion

In a small but highly engaged sample of middle-aged and older adults identifying as B/AA and/or AI/AN, we preliminarily explored relationships between TL and performance on cognitive tests sensitive to age-related cognitive change. Taken as a whole, our findings contribute to the growing body of evidence on biological age as a predictor of cognitive health among Black and Indigenous populations that have been historically underrepresented in and underserved by ADRD research. Descriptive analyses revealed no observable racial differentials in TL within the study sample. While some studies posit that ancestry may account for observable differentials in TL, substantial genetic admixture, geographic variability, and the strong influence of environmental exposures on TL contribute to inconsistencies in this literature (Brown et al., 2017). Further, to our knowledge most or all U.S.-based studies of TL in AI/AN populations have been within-group studies of the Strong Heart Family Study cohort of AI/AN older adults from the Plains and Southwest (Suchy-Dicey et al., 2018; Zhu et al., 2013), with lived experiences and ancestry uniquely distinct from our Great Lakes and Woodland AI/AN cohort.

Our primary analyses suggested that TL may predict later-life cognitive function, specifically immediate memory and learning abilities. Results of nested multivariable regression models indicate that although TL and cognitive performance were partially confounded by chronological age, TL also associated with cognition independently with relationships in the expected direction, where longer telomeres associated with better cognitive test performance scores. While TL estimates were not statistically significant predictors of immediate memory after controlling for chronological age, this may be due to the small sample of participants with data available for this preliminary study. Prior work has empirically established the importance of somewhat larger samples to explore TL as a predictor of health outcomes (Eisenberg et al., 2020). Given the parameter estimates available through our preliminary analysis, we conducted a post-hoc power analysis to provide guidance on minimum sample sizes required for future work, with particular attention to immediate memory as a key outcome among cohorts similar in demographics including age. The estimated sample size needed for adequate power in this domain is well within reach of most cohort studies, assuming strong outreach practices and sufficient time and resource investment in community. Such guidance is particularly relevant for work with communities underrepresented in research but for whom growing and disseminating a community-specific body of evidence is of primary concern and in fact is a key giveback method required for sustained and ethical engagement, recruitment, and retention.

To our knowledge, this is one of relatively few studies of TL and cognitive function among B/AA adults and the only one to intentionally include AI/AN participants. Our study’s evidence of moderate association between TL and immediate memory is inconsistent with findings from at least one study examining similar measures of executive function and episodic memory, who saw no association of TL with test performance in either domain among their similarly sized African American subsample (Leibel et al., 2020), even after allowing relationships to vary by socioeconomic status. However, another longitudinal study with a similarly sized sample reported baseline TL significantly predicted performance in both domains for African American participants (Chen et al., 2023). While our team has not previously examined TL and cognition in the Wisconsin cognitive aging cohorts, we have reported on another socially-rooted “accumulation-of-risk” model, where cumulative life stressors predict of cognitive aging, within both Black and white subsamples (Zuelsdorff et al., 2020). Life stressors are likely to operate on cognition both via short-term, potentially transient mechanisms like depressive symptoms or sleep disruption, and longer-term mechanisms such as biological weathering. Notably, associations between accumulated stressors and executive function were observed for Black and white participants, but associations between accumulated stressors and episodic memory were observed for Black participants only. Our findings for TL in a racially minoritized sample may provide additional support for biological weathering as a contributor to premature changes in memory among these relatively young cohorts.

As noted, a preponderance of studies examining TL as a predictor of cognitive aging have been conducted in predominantly non-Hispanic white samples. Evidence has been mixed in these populations. For instance, though several large studies including meta-analyses report strong linear associations between TL and cognitive outcomes such as risk for dementia (Cao et al., 2023; Hägg et al., 2017; Pudas et al., 2021), other studies report that associations do not withstand adjustment for socioeconomic markers such as educational attainment (Zhan et al., 2018). However, our findings are consistent with those of several other teams. In a sample of middle-aged and older adults drawn from the Spanish Alzheimer’s and Families (ALFA) cognitive aging cohort, TL associated with immediate free recall but not with delayed recall or tests of executive function (Rodríguez-Fernández et al., 2024). Future research on this relationship across the life- and disease course is warranted, especially if longitudinal analysis of interventions to slow biological aging and prevent ADRD are implemented at different life stages.

There are some limitations to this study that should be discussed particularly as they relate to future directions in this crucial area of work. First, small sample sizes not only limited power for whole-sample modeling of direct relationships, but did not allow for exploration of complex relationships that have been observed in larger but predominantly white cohorts. For instance, some studies report that magnitude of effect and even directionality is dependent on individual characteristics such as sex (De Looze et al., 2024; Gampawar et al., 2020) and presence of ADRD pathology in the brain (Mahoney et al., 2019b). Effect modification is not only useful for understanding neurobiological mechanisms and targeting interventions when it is revealed; unmodeled, it may obscure true associations. In the ALFA study cited previously, secondary sex-stratified analyses showed that TL did associate with executive function, but only in females.

We were also unable to look at TL as a time-varying predictor of cognitive health in this cross-sectional pilot study, though such longitudinal associations and evidence for modifiability will be crucial in growing an optimal evidence base for ADRD intervention studies. While ADRD-specific telomere interventions are sparse or non-existent, a meta-analysis of interventions targeting TL found that common interventions combining physical activity with dietary guidance halted telomere attrition and even significantly lengthened telomeres in six to eight weeks (Buttet et al., 2022). Longer-term intervention studies designed to address social and structural determinants of well-being including discrimination-related stress have also reported reduced telomere attrition in treated participants relative to controls (Gilmore-Bykovskyi et al., 2019).

Finally, findings from this volunteer-based cohort cannot be generalized to broader AI/AN and B/AA populations. Cognitive aging study protocols, including those of the source WRAP and ADRC studies, are time- and effort-intensive (Gilmore-Bykovskyi, 2019). As such, they frequently engage highly selected samples with complex motivations for participation that may bias risk profiles. Fortunately, intentional outreach, engagement, and institutional responsivity over time may improve representativeness. Work by our team has found evidence for strong selection bias by enrollment “motivators” such as family history of ADRD and referral from a memory clinic among non-Hispanic white participants in the national ADRC network (Gleason et al., 2019). These correlations were not observed among B/AA participants, who most frequently enroll and share data at sites with strong community engagement and an emphasis on accessibility; however, replication in large population-based cohorts will complement the work done in community-based cognitive aging research.

The current study nonetheless contributes substantially to a small but growing and much-needed body of evidence supporting the use of TL to measure biological age as a modifiable risk factor. If replicated, markers of biological age hold potential as an intervention target and foothold to address change in cognitive domains associated with ADRD among underserved populations at highest risk for developing ADRD. Importantly, the novel findings in this report are the first to demonstrate the relationship between biological age and pereformance in ADRD-specific cognitive domains for the AI/AN population, albeit of a small sample size to date. Targeting preclinical ADRD as well as cognitive changes frorm other neuropathologies will be a major focus in coming decades.

Further, our results strongly suggest that inclusive cohorts, adequately powered for within-group analyses that can replicate and expand on extant findings are within reach for committed research teams. The AA-FAIM study and the Wisconsin ADRC has been successful in collecting biospecimens for AI/AN and B/AA participants due to its culturally appropriate community-based research practices and dedication to inclusive ADRD research (Gleason et al., 2019; Green-Harris et al., 2019). To increase enrollment of participants from minoritized communities, future ADRD research studies should hire underrepresented population community members to the research team, identify culturally appropriate community-based research and dissemination practices specific to the communities of interest, and engage with and respond to populations historically excluded from research through sustained community partnerships.

## Acknowledgements and Funding

The authors humbly thank the Native American Outreach Team and the Black Leaders in Brain Health at the University of Wisconsin ADRC for their support of the creation of the project, suggestions for finding dissemination, and work to increase the diversity of ADRD research. This work was not possible without the meaningful engagement of research participants, their caretakers, families, and communities. This work was supported by several National Institutes of Health awards [U24 AG066528 (Drury); U24 AG066528-S1 (Drury / McLester-Davis); R01 AG054059 (Gleason); P30 AG062715 (Asthana / Johnson / Gleason); P50 AG033514 (Asthana / Johnson); R03 AG063303 (Zuelsdorff)]. Any opinions, findings, and conclusions or recommendations expressed in this material are those of the author(s) and do not necessarily reflect the views of the NIH or the authors’ institutions.

## References

Adkins-Jackson, P. B., George, K. M., Besser, L. M., Hyun, J., Lamar, M., Hill-Jarrett, T. G., Bubu, O. M., Flatt, J. D., Heyn, P. C., Cicero, E. C., Zarina Kraal, A., Pushpalata Zanwar, P., Peterson, R., Kim, B., Turner, R. W., Viswanathan, J., Kulick, E. R., Zuelsdorff, M., Stites, S. D., … Babulal, G. (2023). The structural and social determinants of Alzheimer’s disease related dementias. Alzheimer’s & Dementia: The Journal of the Alzheimer’s Association, 19(7), 3171–3185. 10.1002/alz.13027

Aliberti, M. J. R., Cenzer, I. S., Smith, A. K., Lee, S. J., Yaffe, K., & Covinsky, K. E. (2019). Assessing Risk for Adverse Outcomes in Older Adults: The Need to Include Both Physical Frailty and Cognition. Journal of the American Geriatrics Society, 67(3), 477–483. 10.1111/jgs.15683

Allison, S. L., Jonaitis, E. M., Koscik, R. L., Hermann, B. P., Mueller, K. D., Cary, R. P., Ma, Y., Rowley, H. A., Carlsson, C. M., Asthana, S., Zetterberg, H., Blennow, K., Bendlin, B. B., & Johnson, S. C. (2021). Neurodegeneration, Alzheimer’s disease biomarkers, and longitudinal verbal learning and memory performance in late middle age. Neurobiology of Aging, 102, 151–160. 10.1016/j.neurobiolaging.2021.01.030

Alzheimer’s Association 2024 Alzheimer’s Disease Facts and Figures. (2024) Alzheimers Dement 2024;20(5).

Avila-Rieger, J., Turney, I. C., Vonk, J. M. J., Esie, P., Seblova, D., Weir, V. R., Belsky, D. W., & Manly, J. J. (2022). Socioeconomic Status, Biological Aging, and Memory in a Diverse National Sample of Older US Men and Women. Neurology, 99(19), e2114–e2124. 10.1212/WNL.0000000000201032

Brown, L., Needham, B., & Ailshire, J. (2017). Telomere Length Among Older U.S. Adults: Differences by Race/Ethnicity, Gender, and Age. Journal of Aging and Health, 29(8), 1350–1366. 10.1177/0898264316661390

Bulus, M. (2022). pwrss: Statistical Power and Sample Size Calculation Tools (p. 0.3.1) [Dataset]. 10.32614/CRAN.package.pwrss

Buttet, M., Bagheri, R., Ugbolue, U. C., Laporte, C., Trousselard, M., Benson, A., Bouillon-Minois, J.-B., & Dutheil, F. (2022). Effect of a lifestyle intervention on telomere length: A systematic review and meta-analysis. Mechanisms of Ageing and Development, 206, 111694. 10.1016/j.mad.2022.111694

Byun, M. S., Yi, D., Lee, J. H., Lee, J., Lee, H., Kim, Y. K., Lee, Y., & Lee, D. Y. (2020). Telomere length, brain tau deposition and cognitive decline: Two year follow up study: Biomarkers (non neuroimaging) / Novel biomarkers. Alzheimer’s & Dementia, 16(S4). 10.1002/alz.045919

Canty, A., & Ripley, B. (1999). boot: Bootstrap Functions (Originally by Angelo Canty for S) (p. 1.3-31) [Dataset]. 10.32614/CRAN.package.boot

Cao, Z., Hou, Y., & Xu, C. (2023). Leucocyte telomere length, brain volume and risk of dementia: A prospective cohort study. General Psychiatry, 36(4), e101120. 10.1136/gpsych-2023-101120

Cawthon, R. M. (2009). Telomere length measurement by a novel monochrome multiplex quantitative PCR method. Nucleic Acids Research, 37(3), e21–e21. 10.1093/nar/gkn1027

Chen, C., Yang, K., Nan, H., Unverzagt, F., McClure, L. A., Irvin, M. R., Judd, S., Cushman, M., Kamin Mukaz, D., Klaunig, J. E., D’Alton, M. E., & Kahe, K. (2023). Associations of Telomere Length and Change With Cognitive Decline Were Modified by Sex and Race: The REGARDS Study. American Journal of Alzheimer’s Disease & Other Dementias®, 38, 15333175231175797. 10.1177/15333175231175797

Clark, L. R., Berman, S. E., Norton, D., Koscik, R. L., Jonaitis, E., Blennow, K., Bendlin, B. B., Asthana, S., Johnson, S. C., Zetterberg, H., & Carlsson, C. M. (2018). Age-accelerated cognitive decline in asymptomatic adults with CSF β-amyloid. Neurology, 90(15), e1306–e1315. 10.1212/WNL.0000000000005291

Corrigan, J. D., & Hinkeldey, N. S. (1987). Relationships between Parts A and B of the Trail Making Test. Journal of Clinical Psychology, 43(4), 402–409. 10.1002/1097-4679(198707)43:4<402::AID-JCLP2270430411>3.0.CO;2-E

Davison, A., & Hinkley, D. (1997). Bootstrap Methods and Their Application. Journal of the American Statistical Association, 94. 10.2307/1271471

De Looze, C., McCrory, C., O’Halloran, A., Polidoro, S., Anne Kenny, R., & Feeney, J. (2024). Mind versus body: Perceived stress and biological stress are independently related to cognitive decline. *Brain*, Behavior, and Immunity, 115, 696–704. 10.1016/j.bbi.2023.10.017

Drury, S. S., Mabile, E., Brett, Z. H., Esteves, K., Jones, E., Shirtcliff, E. A., & Theall, K. P. (2014). The Association of Telomere Length With Family Violence and Disruption. Pediatrics, 134(1), e128–e137. 10.1542/peds.2013-3415

Eisenberg, D., Nettle, D., & Verhulst, S. (2020). How to calculate the repeatability (ICC) of telomere length measures.

Elam, K. K., Johnson, S. L., Ruof, A., Eisenberg, D. T. A., Rej, P. H., Sandler, I., & Wolchik, S. (n.d.). Examining the influence of adversity, family contexts, and a family-based intervention on parent and child telomere length. European Journal of Psychotraumatology, 13(1), 2088935. 10.1080/20008198.2022.2088935

Elliott, M. L., Caspi, A., Houts, R. M., Ambler, A., Broadbent, J. M., Hancox, R. J., Harrington, H., Hogan, S., Keenan, R., Knodt, A., Leung, J. H., Melzer, T. R., Purdy, S. C., Ramrakha, S., Richmond-Rakerd, L. S., Righarts, A., Sugden, K., Thomson, W. M., Thorne, P. R., … Moffitt, T. E. (2021). Disparities in the pace of biological aging among midlife adults of the same chronological age have implications for future frailty risk and policy. Nature Aging, 1(3), 295–308. 10.1038/s43587-021-00044-4

Forero, D. A., González-Giraldo, Y., López-Quintero, C., Castro-Vega, L. J., Barreto, G. E., & Perry, G. (2016). Meta-analysis of Telomere Length in Alzheimer’s Disease. The Journals of Gerontology Series A: Biological Sciences and Medical Sciences, 71(8), 1069–1073. 10.1093/gerona/glw053

Gampawar, P., Schmidt, R., & Schmidt, H. (2020). Leukocyte Telomere Length Is Related to Brain Parenchymal Fraction and Attention/Speed in the Elderly: Results of the Austrian Stroke Prevention Study. Frontiers in Psychiatry, 11, 100. 10.3389/fpsyt.2020.00100

Geronimus, A. T. (1992). The weathering hypothesis and the health of African-American women and infants: Evidence and speculations. Ethnicity & Disease, 2(3), 207–221.

Gilmore-Bykovskyi, A., Croff, R., Glover, C. M., Jackson, J. D., Resendez, J., Perez, A., Zuelsdorff, M., Green-Harris, G., & Manly, J. J. (2022). Traversing the Aging Research and Health Equity Divide: Toward Intersectional Frameworks of Research Justice and Participation. The Gerontologist, 62(5), 711–720. 10.1093/geront/gnab107

Gilmore-Bykovskyi, A. L., Jin, Y., Gleason, C., Flowers-Benton, S., Block, L. M., Dilworth-Anderson, P., Barnes, L. L., Shah, M. N., & Zuelsdorff, M. (2019). Recruitment and retention of underrepresented populations in Alzheimer’s disease research: A systematic review. Alzheimer’s & Dementia: Translational Research & Clinical Interventions, 5, 751–770. 10.1016/j.trci.2019.09.018

Gleason, C. E., Martin, W., Strong, L., Summers, M., Lambrou, N. H., Zuelsdorff, M., Chin, N. A., Krainer, J., Lassila, P., Miller, D. A., Jacklin, K., Henderson, J. N., Warry, W., Blind, M. J., Salazar, H., Skenandore, D. W., Edwards, D. F., Carlsson, C., & Asthana, S. (2019). F4-06-02: IMPROVING DEMENTIA OUTCOMES IN INDIAN COUNTRY: THE ONEIDA NATION ALZHEIMER’S DISEASE PROJECT. Alzheimer’s & Dementia, 15, P1227–P1227. 10.1016/j.jalz.2019.06.4734

Glymour, M. M., & Manly, J. J. (2008). Lifecourse social conditions and racial and ethnic patterns of cognitive aging. Neuropsychology Review, 18(3), 223–254. 10.1007/s11065-008-9064-z

Green-Harris, G., Coley, S. L., Koscik, R. L., Norris, N. C., Houston, S. L., Sager, M. A., Johnson, S. C., & Edwards, D. F. (2019). Addressing Disparities in Alzheimer’s Disease and African-American Participation in Research: An Asset-Based Community Development Approach. Frontiers in Aging Neuroscience, 11, 125. 10.3389/fnagi.2019.00125

Grodstein, F., Oijen, M. van, Irizarry, M. C., Rosas, H. D., Hyman, B. T., Growdon, J. H., & Vivo, I. D. (2008). Shorter Telomeres May Mark Early Risk of Dementia: Preliminary Analysis of 62 Participants from the Nurses’ Health Study. PLOS ONE, 3(2), e1590. 10.1371/journal.pone.0001590

Hägg, S., Zhan, Y., Karlsson, R., Gerritsen, L., Ploner, A., van der Lee, S. J., Broer, L., Deelen, J., Marioni, R. E., Wong, A., Lundquist, A., Zhu, G., Hansell, N. K., Sillanpää, E., Fedko, I. O., Amin, N. A., Beekman, M., de Craen, A. J. M., Degerman, S., … Pedersen, N. L. (2017). Short telomere length is associated with impaired cognitive performance in European ancestry cohorts. Translational Psychiatry, 7(4), e1100–e1100. 10.1038/tp.2017.73

Hansen, M. E. B., Hunt, S. C., Stone, R. C., Horvath, K., Herbig, U., Ranciaro, A., Hirbo, J., Beggs, W., Reiner, A. P., Wilson, J. G., Kimura, M., De Vivo, I., Chen, M. M., Kark, J. D., Levy, D., Nyambo, T., Tishkoff, S. A., & Aviv, A. (2016). Shorter telomere length in Europeans than in Africans due to polygenetic adaptation. Human Molecular Genetics, 25(11), 2324–2330. 10.1093/hmg/ddw070

Hochstrasser, T., Marksteiner, J., & Humpel, C. (2012). Telomere length is age-dependent and reduced in monocytes of Alzheimer patients. Experimental Gerontology, 47(2), 160–163. 10.1016/j.exger.2011.11.012

Insel, K. C., Merkle, C. J., Hsiao, C.-P., Vidrine, A. N., & Montgomery, D. W. (2012). Biomarkers for Cognitive Aging Part I: Telomere Length, Blood Pressure and Cognition Among Individuals with Hypertension. Biological Research For Nursing, 14(2), 124–132. 10.1177/1099800411406433

Johnson, S. C., Koscik, R. L., Jonaitis, E. M., Clark, L. R., Mueller, K. D., Berman, S. E., Bendlin, B. B., Engelman, C. D., Okonkwo, O. C., Hogan, K. J., Asthana, S., Carlsson, C. M., Hermann, B. P., & Sager, M. A. (2018). The Wisconsin Registry for Alzheimer’s Prevention: A review of findings and current directions. *Alzheimer’s & Dementia (Amsterdam*, Netherlands), 10, 130–142. 10.1016/j.dadm.2017.11.007

Kennedy, B. K., Berger, S. L., Brunet, A., Campisi, J., Cuervo, A. M., Epel, E. S., Franceschi, C., Lithgow, G. J., Morimoto, R. I., Pessin, J. E., Rando, T. A., Richardson, A., Schadt, E. E., Wyss-Coray, T., & Sierra, F. (2014). Geroscience: Linking aging to chronic disease. Cell, 159(4), 709–713. 10.1016/j.cell.2014.10.039

Kume, K., Kikukawa, M., Hanyu, H., Takata, Y., Umahara, T., Sakurai, H., Kanetaka, H., Ohyashiki, K., Ohyashiki, J. H., & Iwamoto, T. (2012). Telomere length shortening in patients with dementia with Lewy bodies: Telomere shortening in DLB. European Journal of Neurology, 19(6), 905–910. 10.1111/j.1468-1331.2011.03655.x

Lee, M., Whitsel, E., Avery, C., Hughes, T. M., Griswold, M. E., Sedaghat, S., Gottesman, R. F., Mosley, T. H., Heiss, G., & Lutsey, P. L. (2022). Variation in Population Attributable Fraction of Dementia Associated With Potentially Modifiable Risk Factors by Race and Ethnicity in the US. JAMA Network Open, 5(7), e2219672. 10.1001/jamanetworkopen.2022.19672

Leibel, D. K., Shaked, D., Beatty Moody, D. L., Liu, H. B., Weng, N., Evans, M. K., Zonderman, A. B., & Waldstein, S. R. (2020). Telomere length and cognitive function: Differential patterns across sociodemographic groups. Neuropsychology, 34(2), 186–198. 10.1037/neu0000601

Lezak, M. D., Howieson, D. B., Loring, D. W., Hannay, H. J., & Fischer, J. S. (2004). Neuropsychological assessment (4th ed.). Oxford University Press.

Liu, M., Huo, Y. R., Wang, J., Wang, C., Liu, S., Liu, S., Wang, J., & Ji, Y. (2016). Telomere Shortening in Alzheimer’s Disease Patients. Annals of Clinical & Laboratory Science, 46(3), 260–265.

Livingston, G., Huntley, J., Liu, K. Y., Costafreda, S. G., Selbæk, G., Alladi, S., Ames, D., Banerjee, S., Burns, A., Brayne, C., Fox, N. C., Ferri, C. P., Gitlin, L. N., Howard, R., Kales, H. C., Kivimäki, M., Larson, E. B., Nakasujja, N., Rockwood, K., … Mukadam, N. (2024). Dementia prevention, intervention, and care: 2024 report of the Lancet standing Commission. The Lancet, 404(10452), 572–628. 10.1016/S0140-6736(24)01296-0

Lukens, J. N., Van Deerlin, V., Clark, C. M., Xie, S. X., & Johnson, F. B. (2009). Comparisons of telomere lengths in peripheral blood and cerebellum in Alzheimer’s disease. Alzheimer’s & Dementia, 5(6), 463–469. 10.1016/j.jalz.2009.05.666

Mahoney, E. R., Dumitrescu, L., Seto, M., Nudelman, K. N. H., Buckley, R. F., Gifford, K. A., Saykin, A. J., Jefferson, A. J., & Hohman, T. J. (2019). Telomere length associations with cognition depend on Alzheimer’s disease biomarkers. Alzheimer’s & Dementia: Translational Research & Clinical Interventions, 5, 883–890. 10.1016/j.trci.2019.11.003

Mayeda, E. R., Glymour, M. M., Quesenberry, C. P., & Whitmer, R. A. (2016). Inequalities in dementia incidence between six racial and ethnic groups over 14 years. Alzheimer’s & Dementia: The Journal of the Alzheimer’s Association, 12(3), 216–224. 10.1016/j.jalz.2015.12.007

Movérare-Skrtic, S., Johansson, P., Mattsson, N., Hansson, O., Wallin, A., Johansson, J.-O., Zetterberg, H., Blennow, K., & Svensson, J. (2012). Leukocyte Telomere Length (LTL) is reduced in stable mild cognitive impairment but low LTL is not associated with conversion to Alzheimer’s Disease: A pilot study. Experimental Gerontology, 47(2), 179–182. 10.1016/j.exger.2011.12.005

Olson, N. L., & Albensi, B. C. (2020). Race- and Sex-Based Disparities in Alzheimer’s Disease Clinical Trial Enrollment in the United States and Canada: An Indigenous Perspective. Journal of Alzheimer’s Disease Reports, 4(1), 325–344. 10.3233/ADR-200214

Pudas, S., Josefsson, M., Nordin Adolfsson, A., Landfors, M., Kauppi, K., Veng-Taasti, L. M., Hultdin, M., Adolfsson, R., & Degerman, S. (2021). Short Leukocyte Telomeres, But Not Telomere Attrition Rates, Predict Memory Decline in the 20-Year Longitudinal Betula Study. The Journals of Gerontology. Series A, Biological Sciences and Medical Sciences, 76(6), 955–963. 10.1093/gerona/glaa322

R Core Team (2024). _R: A Language and Environment for Statistical Computing_. R Foundation for Statistical Computing, Vienna, Austria. https://www.R-project.org/

Rey Auditory Verbal Learning Test TM: A Handbook. (1996). PAR Inc. from https://www.parinc.com/products/RAVLT

Rodríguez-Fernández, B., Sánchez Benavides, G., Genius, P., Minguillón, C., Fauria, K., De Vivo, I., Navarro i Cuartiellas, A., Molinuevo, J. L., Gispert López, J. D., Sala Vila, A., Vilor Tejedor, N., Crous-Bou, M., & Study, A. (2024). Association between telomere length and cognitive function among cognitively unimpaired individuals at risk of Alzheimer’s disease. 10.1016/j.neurobiolaging.2024.05.015

Scarabino, D., Peconi, M., Broggio, E., Gambina, G., Maggi, E., Armeli, F., Mantuano, E., Morello, M., Corbo, R. M., & Businaro, R. (2020). Relationship between proinflammatory cytokines (Il-1beta, Il-18) and leukocyte telomere length in mild cognitive impairment and Alzheimer’s disease. Experimental Gerontology, 136, 110945. 10.1016/j.exger.2020.110945

Scarabino, D., Broggio, E., Gambina, G., & Corbo, R. M. (2017). Leukocyte telomere length in mild cognitive impairment and Alzheimer’s disease patients. Experimental Gerontology, 98, 143–147. 10.1016/j.exger.2017.08.025

Strauss, E., Sherman, E. M. S., & Spreen, O. (2006). A Compendium of Neuropsychological Tests: Administration, Norms, and Commentary. Oxford University Press.

Suchy-Dicey, A. M., Muller, C. J., Madhyastha, T. M., Shibata, D., Cole, S. A., Zhao, J., Longstreth, W. T., & Buchwald, D. (2018). Telomere Length and Magnetic Resonance Imaging Findings of Vascular Brain Injury and Central Brain Atrophy: The Strong Heart Study. American Journal of Epidemiology, 187(6), 1231–1239. 10.1093/aje/kwx368

Thomas, P., O’ Callaghan, N. J., & Fenech, M. (2008). Telomere length in white blood cells, buccal cells and brain tissue and its variation with ageing and Alzheimer’s disease. Mechanisms of Ageing and Development, 129(4), 183–190. 10.1016/j.mad.2007.12.004

Vaiserman, A., & Krasnienkov, D. (2020). Telomere Length as a Marker of Biological Age: State-of-the-Art, Open Issues, and Future Perspectives. Frontiers in Genetics, 11, 630186. 10.3389/fgene.2020.630186

Wu, J. W., Yaqub, A., Ma, Y., Koudstaal, W., Hofman, A., Ikram, M. A., Ghanbari, M., & Goudsmit, J. (2021). Biological age in healthy elderly predicts aging-related diseases including dementia. Scientific Reports, 11(1), 15929. 10.1038/s41598-021-95425-5

Zhan, Y., Clements, M. S., Roberts, R. O., Vassilaki, M., Druliner, B. R., Boardman, L. A., Petersen, R. C., Reynolds, C. A., Pedersen, N. L., & Hägg, S. (2018). Association of telomere length with general cognitive trajectories: A meta-analysis of four prospective cohort studies. Neurobiology of Aging, 69, 111–116.

Zhu, Y., Voruganti, V. S., Lin, J., Matsuguchi, T., Blackburn, E., Best, L. G., Lee, E. T., MacCluer, J. W., Cole, S. A., & Zhao, J. (2013). QTL mapping of leukocyte telomere length in American Indians: The Strong Heart Family Study. Aging, 5(9), 704–716. 10.18632/aging.100600

Zuelsdorff, M., Okonkwo, O. C., Norton, D., Barnes, L. L., Graham, K. L., Clark, L. R., Wyman, M. F., Benton, S. F., Gee, A., Lambrou, N., Johnson, S. C., & Gleason, C. E. (2020). Stressful Life Events and Racial Disparities in Cognition Among Middle-Aged and Older Adults. Journal of Alzheimer’s Disease, 73(2), 671–682. 10.3233/jad-190439

